# A Forward Genetic Screen in *Caenorhabditis elegans* for Genes that Modulate α-synuclein-Induced Neurodegeneration through the Mitochondrial UPR

**DOI:** 10.64898/2026.06.11.731199

**Authors:** Karolina Willicott, Joy D. Iroegbu, Madeline R. Greene, Aidan C. Meyers, Rachel Martin, Rose Davidson-Tullis, Laura A. Berkowitz, Guy A. Caldwell, Kim A. Caldwell

## Abstract

Overexpression of α-synuclein (α-syn), an inherently disordered protein, triggers chronic activation of the mitochondrial unfolded protein response (UPR^mt^) pathway in *Caenorhabditis elegans* with enhanced dopaminergic (DAergic) neurodegeneration. Introduction of a loss-of-function*(lf)* mutation in *atfs-1*, the main transcriptional regulator of the UPR^mt^, into α-syn nematodes results in significant neuroprotection from α-syn-induced DA neuron loss, indicating that compensatory mechanisms provide neuroprotection. We performed a F3 forward genetic screen in *C. elegans atfs-1(lf)* mutants to identify molecular components associated with the modulation of neurodegeneration via UPR^mt^ signaling in α-syn-expressing DA neurons. Homozygous mutant animals were examined for enhanced neurodegeneration; multiple independent alleles were uncovered. Among these, we identified new nonsense alleles encoding the histone lysine demethylases (H3K27me3), *jmjd-1.2* and *jmjd-3.1*. Another line carried a nonsense allele of *twk-14.* This gene encodes a conserved protein termed KCNK12 in mammals that facilitates passive background K^+^ leak currents to set and stabilize resting membrane potential. To further examine the association of these gene products in DA neurodegeneration, mutants and/or RNA interference were employed. DA neurodegeneration was observed in the α-syn *+ atfs-1(lf)* background when *jmjd-1.2*, *jmjd-3.1,* or *twk-14* were individually depleted. These results provide evidence that *jmjd-1.2* and *jmjd-3.1*, which encode previously characterized H3K27me3 demethylases, and the uncharacterized *twk-14* gene product, orthologous to human KCNK12, naturally confer protection from α-syn neurotoxicity.

## Introduction

Among the unresolved molecular mechanisms associated with Parkinson disease (PD) are mitochondrial dysfunction and the neurodegenerative effects related to the accumulation of misfolded α-synuclein (α-syn) protein. Previously, α-syn was shown to interact with TOMM20, an outer mitochondrial membrane protein (Di Maio et al., 2016). The cellular consequences of this interaction included impaired protein import into the mitochondrial matrix, reduced respiration, and enhanced generation of reactive oxygen species (ROS). Normally, approximately 1,500 nuclear-encoded proteins pass through the translocase of the outer membrane receptors and continue through the inner membrane into the mitochondrial matrix (Lionaki and Tavernarakis, 2013). If these proteins are hindered from entering the matrix, mitochondrial stress will result (Baker and Haynes, 2011).

Cells monitor protein import efficiency to gauge mitochondrial function. Surveillance mechanisms that recognize aberrant accumulation of proteins destined for mitochondrial import and ROS levels to integrate changes in cytosolic proteostasis with nuclear response (Sutanday et al., 2023). Under stress conditions, the mitochondrial unfolded protein response (UPR^mt^) pathway is activated. A main transcriptional regulator of this pathway is the ATFS-1 protein (<u>A</u>ctivating <u>T</u>ranscription <u>F</u>actor associated with <u>S</u>tress <u>1</u>). We previously determined that the UPR^mt^ is significantly triggered in response to transgenic multicopy human α-syn expression in *C. elegans*, where it is expressed persistently in dopaminergic (DA) neurons when driven by the P*_dat-1_* promoter (Martinez et al., 2017). Moreover, induction of ATFS-1 activity enhances α-syn-induced DA neurodegeneration in transgenic *C. elegans.* Strikingly, when *atfs-1* is deleted, α-syn-induced DA neurodegeneration is robustly suppressed (Figure 1A) (Martinez et al., 2017). We hypothesized that this genetic background would provide an opportunity to identify compensatory molecules that modulate the consequences of aberrant UPR^mt^ induction through a classical forward genetic suppressor screen.

**Figure 1.**
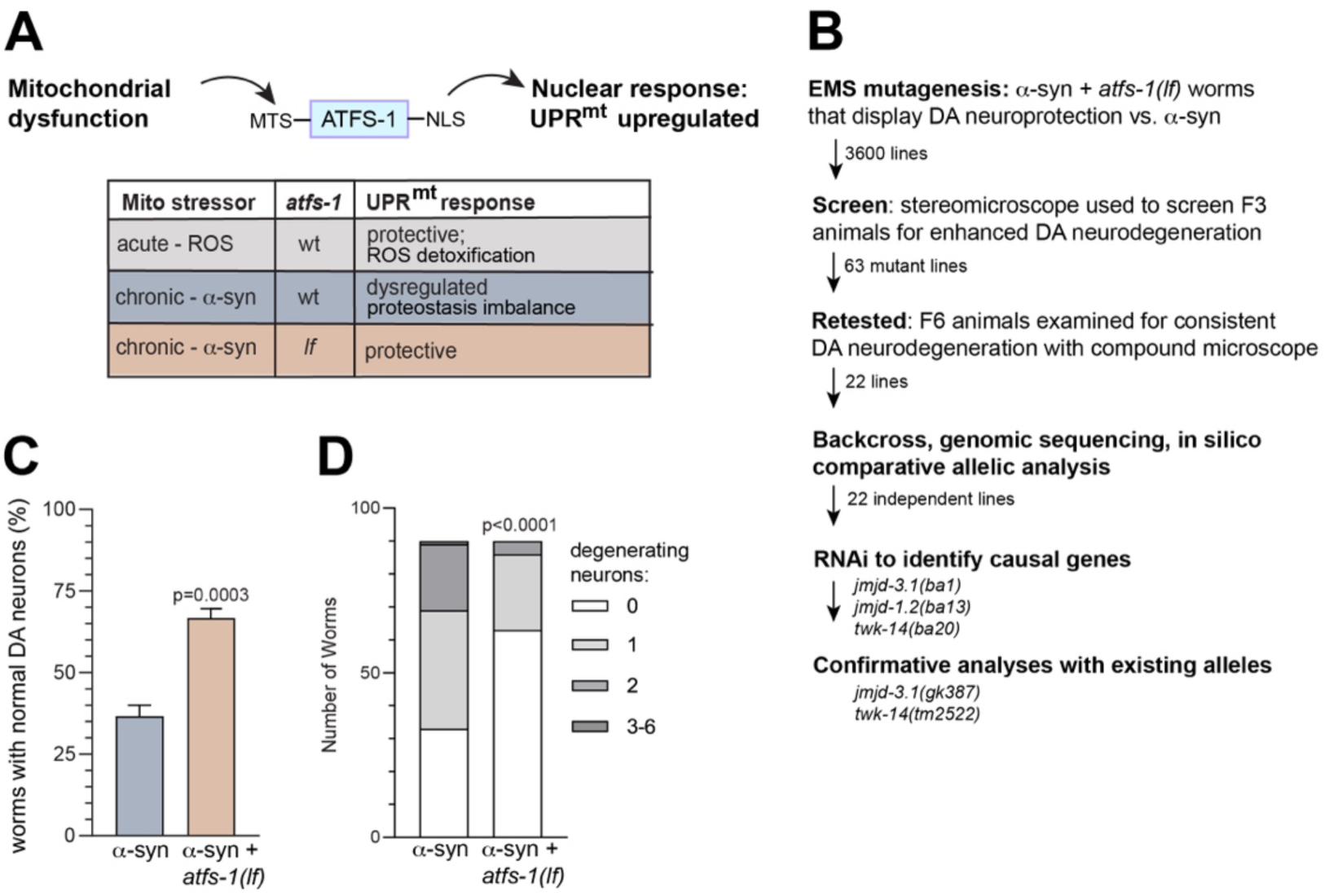
Reduction in α-syn-induced neurodegeneration in the *atfs-1* loss-of-function background establishes a phenotype for identifying mutants that modulate this phenotype. (A) Broad overview of the relationship between ATFS-1 and the UPR^mt^. Via its mitochondrial targeting sequence (MTS), ATFS-1 typically localizes to mitochondria, where it is degraded until stress induces relocalization to the nucleus via a nuclear localization sequence (NLS) to transcribe hundreds of cytoprotective genes, collectively termed the UPR^mt^ (B) Design of the genetic screen and results. Three thousand six hundred F2 lines were established following EMS mutagenesis. The F3 progeny were screened for enhanced DA neurodegeneration using stereomicroscopy. Sixty-three lines were rescreened for neurodegeneration using compound microscopy at the F5 generation as a secondary screen; 22 lines displayed enhanced DA neurodegeneration. These lines were backcrossed three times; their genomic DNA was then sequenced and *in silico* comparative allelic analysis was performed to prioritize candidate targets. RNAi gene knockdown was performed on gene candidates of nonsense mutations using non-EMS treated α-syn; *atfs-1(lf)* worms. Three targeted genes corresponding to putative mutant alleles resulted in enhanced α-syn-induced DA neurodegeneration by RNAi knockdown. Further experimentation confirmed causal mutations and upregulation of the UPR^mt^ via these mutations. (C) Overexpression of α-synuclein (α-syn) under control of a DAergic promoter (P*_dat-1_*::α-syn) results in DA neurodegeneration. Neurodegeneration is readily scored in individual DA neurons using a compound fluorescent microscope in worms that also separately express GFP (P*_dat-1_*::GFP) using a magnification of 600X. When a loss-of-function*(lf)* mutation in *atfs-1(gk3094)* is introduced into these worms [P*_dat-1_*:: α-syn; P*_dat-1_*::GFP; *atfs-1(lf)*], DA neurodegeneration is significantly decreased. N=3; n=30; Student’s two-tailed type 2 *t*-test (D) Individual neuron counts for each strain (from C) displaying the distribution of total neurons categorized as degenerative. Chi square test was performed comparing α-syn; *atfs-1(lf)* to the α-syn alone control. N = 3; n = 30; *p* < 0.0001.

*C. elegans* is experimentally amenable for genetic dissection of mechanisms associated with the consequences of the UPR^mt^ in DA neurons. Results of prior forward genetic screens in *C. elegans* identified alleles associated with DA neuron differentiation (Doitsidou et al., 2008), dopamine signaling mutants using a behavioral assay (Hardaway et al., 2012), and alleles that were protective against 6-hydroxydopamine toxicity in DA neurons (Nass et al., 2005; Masoudi et al., 2014).

Here, we describe the first forward genetic screen to identify genes that contribute to UPR^mt^-associated neuroprotection using *C. elegans* expressing α-syn in DA neurons. From multiple mutagenized lines that enhanced neurodegeneration in worms expressing α-syn in a genetic background lacking functional ATFS-1 [α-syn + *atfs-1(lf)*], we successfully identified and characterized three alleles. New nonsense alleles in the histone lysine (H3K27me3) demethylases *jmjd-1.2* and *jmjd-3.1* were uncovered (Figure 1B). These H3K27me3 demethylases have previously characterized roles in UPR^mt^ activation (Merkwirth *et al*. 2016; Tian *et al*. 2016). To confirm our findings, we determined that RNAi depletion of both *jmjd-1.2* and *jmjd-3.1* enhanced α-syn-induced neurodegeneration in *C. elegans* DA neurons in the *atfs-1(lf)* mutant background. An independent *jmjd-3.1* deletion mutant was also crossed into *C. elegans* expressing α-syn that were either *atfs-1* wildtype or *atfs-1(lf)* mutant and the loss of *atfs-1* was significantly more neuroprotective.

An additional new modifier was identified as a nonsense allele of *twk-14*, an ortholog of mammalian *KCNK12*. The *KCNK12* gene product encodes a two-pore domain potassium leak (K2P) channel. Most research on KCNK12 has focused on cellular localization and/or heterologous expression studies of ion channel activity (Aggarwal *et al*., 2021). This newly identified *twk-14* allele significantly enhanced α-syn-induced neurodegeneration. In a background without α-syn expression, there was no evidence of DA neurodegeneration, thereby uncovering a novel *in vivo* function for KCNK12-like channels that is dependent on α-syn. Furthermore, the genetic combination of *atfs-1(lf)* and *twk-14* mutants exhibited robust neurodegeneration in response to α-syn overexpression.

Taken together, the outcomes of this genetic screen provide a window through which the UPR^mt^ can be viewed in the biomedical context of neurodegenerative disease. These newfound mutations extend understanding of the functional consequences of disrupting JMJD-1.2/JMJD-3.1-associated transcriptional regulation when the UPR^mt^ is chronically activated. Furthermore, the identification of a *twk-14* mutant as an effector of α-syn neurotoxicity places TWK-14/KCNK12 activity at the center of pathogenic factors modifying DA neuron vulnerability and survival in PD and related disorders.

## Materials and Methods

### *C. elegans* culture and strains

Strains were maintained at 20°C on nematode growth media (NGM) agar plates seeded with *E. coli* OP50 bacterial lawn as specified by Brenner (1974). Strains used in this study are listed in Table 1. The entire list of alleles identified in the screen are itemized in Table S1.

**Table 1.** Strains used in this study.

| Strain name | Genotype |
| --- | --- |
| BY250 | [*vtIs7*] *P<sub>dat-1</sub>::GFP* |
| PHX6863 | *twk-14(syb6863)* |
| VC936 | *jmjd-3.1(gk387)* |
| UA44 | [*bals11*] *P<sub>dat-1</sub>::α-syn*; *P<sub>dat-1</sub>::GFP* |
| UA196 | *sid-1(pk3321)*; [*bals11*] *P<sub>dat-1</sub>::α-syn*; *P<sub>dat-1</sub>::GFP*; [*bals33*] *P<sub>dat-1</sub>::sid-1*; *P<sub>myo-2</sub>::mCherry* |
| UA460 | *atfs-1(gk3094)*; [*bals11*] *P<sub>dat-1</sub>::α-syn*; *P<sub>dat-1</sub>::GFP* |
| UA469 | *atfs-1(gk3094)*; *sid-1(pk3321)*; [*bals11*] *P<sub>dat-1</sub>::α-syn*; *P<sub>dat-1</sub>::GFP*; [*bals33*] *P<sub>dat-1</sub>::sid-1*; *P<sub>myo-2</sub>::mCherry* |
| UA475 | *jmjd-3.1(gk387)*; *atfs-1(gk3094)*; [*bals11*] *P<sub>dat-1</sub>::α-syn*; *P<sub>dat-1</sub>::GFP* |
| UA476 | *jmjd-3.1(gk387)*; [*bals11*] *P<sub>dat-1</sub>::α-syn*; *P<sub>dat-1</sub>::GFP* |
| UA477 | *twk-14(syb6863)*; [*vtIs7*] *P<sub>dat-1</sub>::GFP* |
| UA478 | *twk-14(syb6863)*; [*bals11*] *P<sub>dat-1</sub>::α-syn*; *P<sub>dat-1</sub>::GFP* |
| UA484 | *jmjd-3.1(ba5)*; *atfs-1(gk3094)*; [*bals11*] *P<sub>dat-1</sub>::α-syn*; *P<sub>dat-1</sub>::GFP* |
| UA492 | *jmjd-1.2(ba13)*; *atfs-1(gk3094)*; [*bals11*] *P<sub>dat-1</sub>::α-syn*; *P<sub>dat-1</sub>::GFP* |
| UA499 | *twk-14(ba20)*; *atfs-1(gk3094)*; [*bals11*] *P<sub>dat-1</sub>::α-syn*; *P<sub>dat-1</sub>::GFP* |

### Establishing screening conditions

Plates containing 20-30 UA44 (P*_dat-1_*::α-syn; P*_dat-1_*::GFP) and UA460 [P*_dat-1_*::α-syn; P*_dat-1_*::GFP; *atfs-1*(*gk3094*)] worms that had been grown until day 7 at 20°C were placed under an SMT1 fluorescent stereomicroscope (Tritech Research, Los Angeles, CA) and blindly assessed for illuminated cell body numbers using a GFP long pass filter cube (450-490nm ex; 510nm+ em). Using 25X magnification, the cell bodies of GFP-labeled anterior DA neurons (4 CEP and 2 ADE) within individual worms were scored from 0 (no visible cell bodies) to 6 (all cell bodies present). Two plates/strain of worms were scored (>20 worms/plate/strain); the average number of cell bodies per worm was calculated for each plate. The values from two plates/genetic background were averaged to compute an anterior DA neuron score using a Student’s two-tailed type 2 *t*-test (GraphPad Prism, version 10.2.3).

### EMS Mutagenesis

UA460 worms [P*_dat-1_*::α-syn; P*_dat-1_*::GFP; *atfs-1*(*gk3094*)] were grown at 20°C until animals reached the L4 stage (62 hours post 3-hour egg-lay) using standard methods (Brenner, 1974). These animals were mutagenized with 0.05 M ethyl methanesulfonate (EMS) (Sigma #M-0880) as previously described (Kutscher and Shaham, 2014). Briefly, mutagenized worms were placed onto a 10 cm NGM plate and allowed to recover for 4 hours before being transferred to a new plate that did not have EMS. The P0 animals were allowed to lay eggs overnight and then 200 F1 adults were individually transferred to new plates. Four F2 animals from each plate were cloned onto new plates and the F3 progeny were scored at day 7 post-hatching using a stereo fluorescent microscope to identify lines of worms displaying substantial DA neuron cell death, as described in more detail in the screen section below.

### Screen: suppression of α-syn + *atfs-1(lf)* neuroprotection

Scoring F3 animals consisted of a brief, yet inclusive, examination of all worms on each agar plate of cloned, EMS-mutagenized animals that were grown until day 7 at 20°C. An SMT1 fluorescent stereomicroscope (Tritech Research, Los Angeles, CA) was used at the 25X setting to identify lines of UA460 animals for a suppressor phenotype indicative of enhanced DA neuron survival. Mutagenized lines were retained if most animals displayed ≤ 3 DA neuronal cell bodies (out of 6 anterior DA neuronal cell bodies) on average. To ensure homozygosity of candidate mutant worm strains, up to 10 individual worms from each mutant line were cloned onto new plates and subsequently grown for two more generations. At the F5 generation, cell body loss was screened again at the stereomicroscope level. Homozygous animals displaying the suppressor phenotype were then retested using compound fluorescent microscopy at the F6 generation.

### Retest: Neurodegeneration analysis of homozygous mutants

The DA neurons from positive candidates (F6 generation) were examined for neurodegeneration at the compound fluorescent microscope level using the Nikon Eclipse E600 (Tokyo, Japan). Synchronous populations of worms were achieved by egg laying three replicates of 10-15 animals for 3 hours. Worms were grown until day 4 at 20°C and then transferred every day until analysis at day 7. For each strain, 30 animals were mounted on agarose slides within a drop of 1 μM levamisole. Each strain was analyzed in triplicate, for a total of 90 animals at magnification of 600X. The six anterior DA neurons were assessed for neurodegeneration as described previously (Berkowitz *et al*., 2008). Briefly, criteria for DA neurodegeneration included cell body loss, neurons that were excessively dim, completely missing, had broken projections, or projections that did not extend to full normal length. An animal was scored as normal if it possessed all six anterior DA neurons that did not display signs of degeneration. Statistical analysis was performed by comparing each mutant to the UA460 control [α-syn + *atfs-1(lf)*] using the Student’s two-tailed type 2 *t*-test (GraphPad Prism v10.2.3).

### Outcrossing of mutant strains

Only mutants that were significantly more neurodegenerative than the UA460 control were maintained and backcrossed. Ten unmutagenized UA460 males were crossed with three EMS mutagenized hermaphrodites. Ten male offspring from the first outcross were mated to a fresh stock of unmutagenized UA460 hermaphrodites during the second outcross. This was repeated once more for a total of three backcrosses per strain. After the third backcross, ten hermaphrodite offspring per mutagenized line were cloned onto individual plates and their progeny examined for evidence of DA neurodegeneration on day 7. Backcrossed lines that were significantly neurodegenerative compared to UA460 under a compound fluorescent microscope were considered homozygous and were maintained for further experimentation.

### DNA extraction, library preparation, and genome sequencing

To grow sufficient mutant worms for genomic DNA extraction, 70 worms of each mutant line were placed onto five 10 cm NGM plates seeded with *E. coli* OP50 and allowed to lay eggs for 24 hours before removal. The offspring were grown until they reached the adult stage. Genomic DNA was extracted using the Gentra PureGene Kit (QIAGEN Inc, Germantown, MD) following instructions from the manufacturer. DNA was then sent to Novogene Corporation, Inc (Sacramento, CA) for whole genome sequencing. An Illumina NovaSeq 6000 platform was used to obtain short reads of all samples. Another 70 worms of the control strain UA460 were grown as described above, for creation of a reference genome. High molecular weight DNA for this reference genome was extracted using Nanobind Nematode Big DNA Protocol (Circulomics) for Oxford NanoPore (ONT) sequencing. The DNA library was prepared with the ONT Ligation Sequencing Kit (SQK-LSK110) using instructions from the manufacturer (https://community.nanoporetech.com/protocols/genomic-dna-by-ligation-sqk-lsk110/checklist_example.pdf?device=minion). A total of 1.2 μg was used to construct the DNA library in the flow cell. GridION sequencing was performed using a R9.4.1 SpotON flow cell (FLO-MIN106D), using the default script from the MinKNOW program, and files were obtained in the FASTQ format.

### Bioinformatic analysis and in silico comparative allelic analysis

Raw fastq reads were generated by the Guppy basecaller within MiniKNOW. Reads were concatenated into one file, and adapters were trimmed using Porechop with the parameter “--discard_middle”. A preliminary assembly was generated using Canu (version 2.1). Flye (version 2.8.3) was used with the parameter “--nano-corr” to further correct and assemble the genome. Illumina short reads were then used to polish the assembly using Pilon (version 1.23). Four rounds of polishing were done to achieve the most accurate identification of genes (Sutton *et al*., 2021). After assembly, BLAST (version 2.9.0) was used to check the genome for contamination. For annotation, the assembly was first scaffolded using Ragtag (version 2.0.1), then annotated with Liftoff (version 1.6.1). SNPs from the mutant strains were identified using the Genome Analysis Toolkit (GATK) (version 3.4-0-g7e26428). Once SNPs were identified by GATK, genes common to multiple datasets were identified using a custom Python script implemented with pandas (McKinney, 2010). The script is available on GitHub (https://github.com/aidanmeyers/gene-merger-script). Gene lists from individual datasets were merged and grouped by gene name, and the frequency of occurrence of each gene across datasets was determined. Genes present in more than one dataset were retained, and associated annotations were consolidated into a summary table for downstream analysis.). Gene lists from individual datasets were merged and grouped by gene name, and the frequency of occurrence of each gene across datasets was determined. Genes present in more than one dataset were retained, and associated annotations were consolidated into a summary table for downstream analysis.

### RNA interference

Bacterial RNAi clones targeting knockdown of candidate genes, including *twk-14(K01D12.4), jmjd-1.2(F29B9.2)* and *jmjd-3.1(F18E9.5)* were obtained from the Ahringer RNAi Library (Kamath and Ahringer, 2003). *E. coli* (HT115) containing each RNAi feeding vector were initially grown on LB plates that included 12.5 µg/mL tetracycline and 100 µg/mL ampicillin for 16 hours at 37°C. Single colonies were then transferred to LB broth containing 100 µg/mL ampicillin for 16 hours at 37°C. RNAi plates, which also contained 1mM IPTG and 100 µg/mL ampicillin, were seeded with 100 µL of RNAi bacteria and grown for 24 hours at ambient temperature. *C. elegans* strain UA469 [P*_dat-1_*::α-syn; P*_dat-1_*::GFP; *sid-1*(*pk3321*); P*_dat-1_*::*sid-1*; P*_myo-2_*::mCherry; *atfs-1*(*gk3094*)] was exposed to the RNAi bacteria and then scored for neurodegeneration on day 7 as described previously (Hamamichi et al., 2008).

### Sanger sequencing confirmation of mutants

To extract gDNA for Sanger sequencing, 5-7 young adults from *C. elegans* strains UA480 [*jmjd-3.1*(*ba5*); P*_dat-1_*::α-syn; P*_dat-1_*::GFP; *atfs-1*(*gk3094*)], UA492 [*jmjd-1.2*(*ba13*); P*_dat-1_*::α-syn; P*_dat-1_*::GFP; *atfs-1*(*gk3094*)], and UA499 [*twk-14*(*ba20*); P*_dat-1_*::α-syn; P*_dat-1_*::GFP; *atfs-1*(*gk3094*)] were placed into a PCR tube containing 10μL of water and frozen in liquid nitrogen. A master mix was prepared consisting of 15μL of Worm Lysis Buffer [50mM KCl, 10mM Tris (pH 8.3), 2.5mM MgCl_2_, 0.45% NP-40, 0.45% Tween-20, 0.01% gelatin, water] and 0.09μL of Proteinase K [NEB molecular grade (P-8107S)], per sample, was distributed to each still-frozen tube. Tubes were then placed into a thermocycler and incubated at 50°C for 60 minutes and 99°C for 15 minutes. The extracted DNA was processed for Sanger sequencing (Eurofins Genomics; Louisville, KY). After sequencing, Clustal Omega Multiple Sequence Alignment tool (Madeira *et al*., 2022) was used to align mutant sequences to each of the reference gene sequences sourced from Wormbase (wormbase.org).

### Creation of CRISPR *twk-14* strain

Genome editing of *C. elegans* (wildtype, N2) was performed to generate the *twk-14*(*syb6863*) mutant allele, strain PHX6863, using the CRISPR technique by SunyBiotech (Fujian, China). The specific sgRNAs used in creating the strain were:

Sg1: ATGTGTCTTCAATTGTTGTTAGG

Sg2: GAATTGGATGATCAAGAAAATGG

Synonymous mutations were introduced into the PAM sites. *C. elegans* strain PHX6863 contains a single nucleotide polymorphism in the *twk-14* gene encoding a W358STOP change in the TWK-14 protein. The *twk-14*(*syb6863*) mutant was outcrossed twice and then genetically crossed to BY250 {[*vtIs7*] P*_dat-1_*::GFP} and UA44 {[*baIs11*] P*_dat-1_*::α-syn; P*_dat-1_*::GFP} to yield strains UA477 and UA478, respectively. The genetic lesion was confirmed using PCR and allele-specific restriction digestion with *DraI*; the *twk-14* mutant form yields a fragment while a WT *twk-14* does not.

### Neurodegeneration analysis of candidate mutant and CRISPR-modified animals

Confirmation of neurodegenerative phenotypes after final outcrossing and RNAi validation was performed for the three candidate mutant strains UA480, UA492, UA499, as well as the final CRISPR-modified animals (UA478). DA neurodegeneration was quantified as described above (in “Retest: Neurodegeneration analysis of homozygous mutants”) at the compound fluorescent microscope level. Statistical analysis was again performed by comparison to the UA460 control (for candidate mutants) or UA477(CRISPR-modified worms) using the Student’s *t*-test p<0.05 (GraphPad Prism v10.2.3). Individual neuron data collected were also reported in some of the graphs as a distribution from within an entire pool of neurons for each strain examined. Degenerating DA neurons (0 through 6) were analyzed using a Chi square test and compared to control.

### Protein modeling

Three-dimensional structural models of both the wildtype and nonsense mutant proteins encoded by jmd-1.2, jmd-3.1, and twk-14 were generated using the AlphaFold 3 server (Abramson et al., 2024). The resulting predicted structures were imported into UCSF ChimeraX v.1.8 for visualization and comparative analysis (Pettersen et al., 2021). To assess structural variations, each nonsense mutant was superimposed onto its respective wild-type counterpart using the Matchmaker algorithm within ChimeraX. For visualization purposes, the wildtype and nonsense structures of *twk-14* are presented as a single superimposed overlay to highlight their overlapping topologies. Conversely, to ensure visual clarity of the broader conformational changes, the aligned wildtype and mutant structures of *jmjd-3.1* and *jmjd-1.2* are displayed in separate images.

### Data availability

*C. elegans* strains are available upon request. Scripts for data analysis can be accessed at: https://github.com/aidanmeyers/gene-merger-script Whole genome sequencing data have been deposited in the NCBI Sequence Read Archive; the project can be viewed via accession number PRNJA1476676. File S1 describes the contents of the supplemental files.

## Results and Discussion

### Overview of genetic screen

Our classical forward genetic screen (Figure 1B) was designed to take advantage of a previous observation where we found that *C. elegans* overexpressing human α-syn, specifically in DA neurons illuminated by GFP (P*_dat-1_*::α-syn; P*_dat-1_*::GFP), displayed significant neuroprotection when crossed with *atfs-1(gk3094)*, a deletion mutation (Figure 1C, Figure 1D). The 881 base pair deletion in *atfs-1(gk3094)* begins in exon 3, which is predicted to result in a frameshift affecting the essential leucine zipper and nuclear localization signal domains (Wu *et al*., 2018); this causes a loss-of-function [*atfs-1(lf)*] (Martinez *et al*., 2017; Soo *et al*., 2021). As schematized in Figure 1A, the UPR^mt^ is most often characterized as a cytoprotective pathway that is triggered to manage acute stressors, such as toxins (Durieux *et al*., 2011; Rauthan *et al*., 2013; Pellegrino *et al*., 2014). However, we determined that in the presence of α-syn, a chronic UPR^mt^ stressor, dysregulation occurs (Martinez *et al*., 2017). ATFS-1 is the key regulator to promote mitochondrial proteostasis via the UPR^mt^. We reasoned that in our animals, which display significant neuroprotection from α-syn-induced DA neuron loss in an *atfs-1(lf)* background, an opportunity existed to identify supporting molecules associated with this process via a forward classical genetic screen (Figure 1B).

### Establishing screening conditions

There is significant DA neurodegeneration when worms overexpress α-syn specifically within the DA neurons [UA44 (P*_dat-1_*::α-syn; P*_dat-1_*::GFP)]. In Figure 1C, this is displayed as worm population data. Here, an α-syn-expressing animal is considered abnormal if at least one neuron per animal displays a degenerative change. Additionally, DA neurodegenerative changes for all the neurons within a population of the same animals was also depicted as neuron counts representing the severity of DA neuron degeneration as a distribution (0 through 6) for each strain examined. (Figure 1D). Significant neuroprotection from α-syn-induced DA neuron loss was observed in the *atfs-1(lf)* background [UA460, *atfs-1*(*lf*); (P*_dat-1_*::α-syn; P*_dat-1_*::GFP)] (Figure 1, C and D). These phenotypic data were methodically gathered using a compound fluorescent microscope. However, we were curious to know if these differences could be readily distinguished using stereo fluorescent microscopy, a platform far more amenable to genetic screening.

To establish our screening conditions using a stereo fluorescent microscope, we initially asked if we could observe neurodegeneration differences between α-syn (UA44) and α-syn + *atfs-1*(*lf*) (UA460) worms at much lower magnification. Using a 25X setting on a stereo fluorescent microscope, the cell bodies (only) of GFP-labeled DA neurons within individual worms were readily distinguishable while the neurites were no longer visible, as they were when viewing our animals with a compound fluorescent microscope. Moreover, at this lower magnification, multiple worms were observed simultaneously, allowing for inter-animal comparison within a strain. We blindly scored the number of cell bodies in α-syn and α-syn + *atfs-1*(*lf*) *C. elegans*. A rating system was based on the total possible number of anterior DA neurons. Within an individual worm, a score of 6 denoted that all anterior neuron cell bodies were visible [2 ADE (anterior deirid); 4 CEP (cephalic sensory)] while a 0 indicated that no cell bodies were discernible. We determined there was a significant difference between the two strains. On average, the α-syn + *atfs-1*(*lf*) animals displayed 4.6 anterior DA neurons at the stereo fluorescent microscope, while in worms overexpressing α-syn with wildtype *atfs-1* this number was 3.5 neurons (p=0.0004) (Figure 2A).

**Figure 2.**
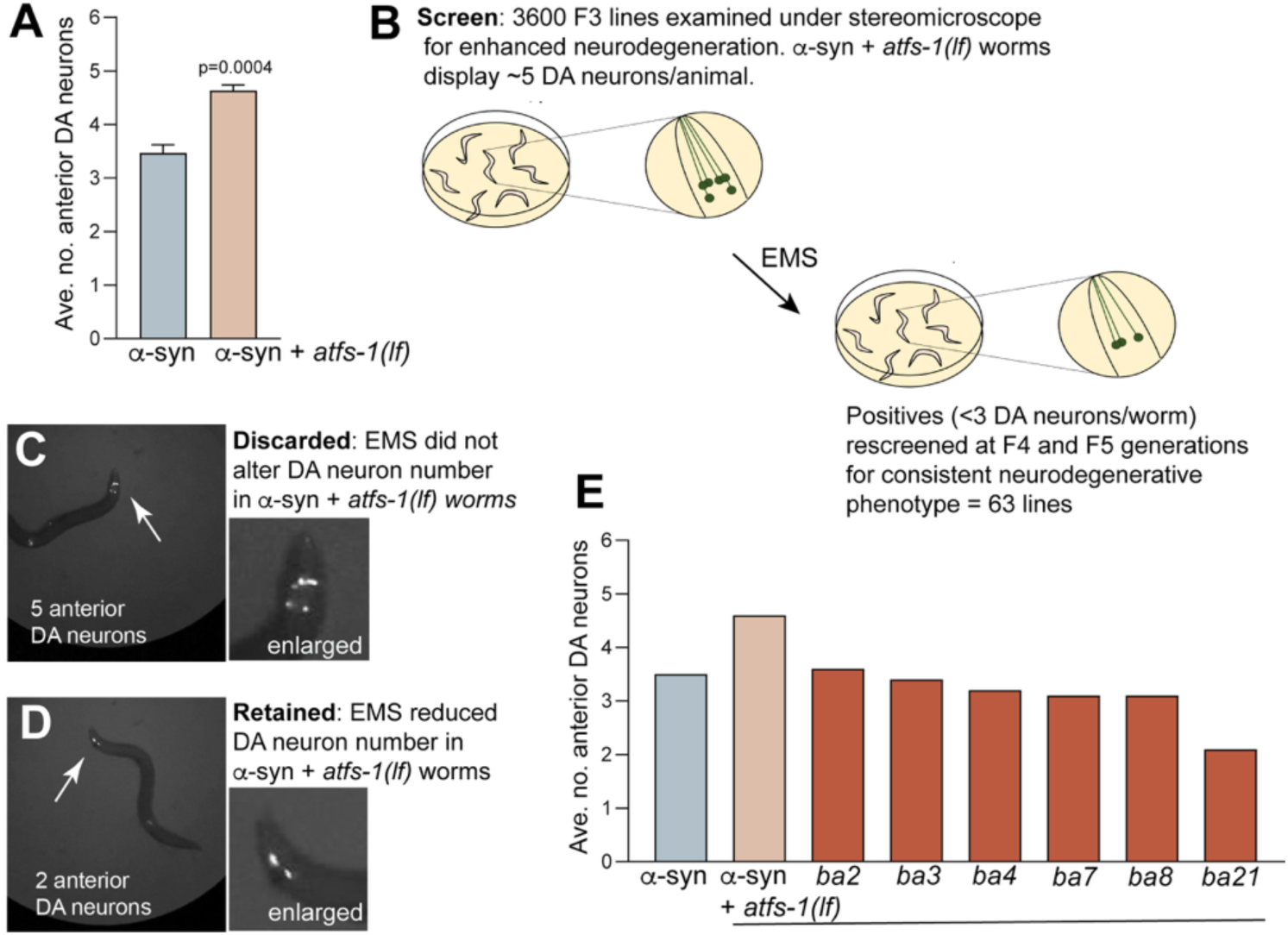
Screen for suppression of DA neuroprotection in α-syn + *atfs-1(lf)* worms using a stereo fluorescent microscope. (A) The cell bodies of GFP-labeled anterior DA neurons from 20-30 α-syn (UA44) and α-syn + *atfs-1(lf)* (UA460) worms were scored from 0 (no visible cell bodies) to 6 (all cell bodies present) using 25X magnification. Screening was performed on day 7, post-hatching. The values from two plates/genetic background were averaged to compute an anterior DA neuron score using a Student’s two-tailed type 2 *t*-test. (B) Diagram of the suppression screen. Cell body neurodegeneration was assessed for each mutagenized line. (C, D) Exemplar images of DA cell bodies illuminated by GFP in α-syn + *atfs-1(lf)* worms [full genotype: [*atfs-1*(*gk3094*); P*_dat-1_*::α-syn; P*_dat-1_*::GFP] before (C) and after (D) treatment with EMS. In image C, there are 5/6 anterior DA neurons, reflective of non-mutagenized, α-syn + *atfs-1(lf)* animals. In image D, only 2/6 anterior DA neurons remain, indicating a possible suppressor phenotype. (E) Example mutant lines identified from the screen (*ba2 – ba4, ba7, ba8, ba21*) displaying suppression of the DA neuroprotection observed in α-syn + *atfs-1(lf)* compared to α-syn alone.

### Screen: suppression of α-syn + *atfs-1(lf)* neuroprotection

We established significant DA cell body differences between α-syn and α-syn + *atfs-1(lf) C. elegans* were readily visible at the stereomicroscope level. Therefore, the α-syn + *atfs-1(lf)* worms were mutagenized with EMS and F2 lines established, which were used to create F3 progeny. At day 7, the F3 α-syn + *atfs-1(lf)* progeny were screened for the DA neurodegeneration suppressor phenotype. In total, ∼3,600 genomes were screened at the F3 generation (Figure 1B and Figure 2B). Control, unmutagenized α-syn worms consistently displayed 1 DA neuron less, on average, than did α-syn + *atfs-1*(*lf*) animals (Figure 2A). Therefore, while screening the latter worm strain in the suppressor screen, if most worms on a plate possessed ≥4 anterior DA cell bodies, the plate was discarded (Figure 2B and 2C), while if the worms on a plate exhibited ≤3 DA cell bodies on average, the plate of animals was retained (Figure 2B and 2D).

To ensure homozygosity of candidate mutant worm strains, up to 10 individual worms from each plate were cloned onto new plates to establish new lines and grown for two more generations (Figure 2B). At the F5 generation, cell body loss was once again screened at the stereomicroscope level. Examples of lines that retained the suppressor phenotype through the F5 generation were displayed relative to unmutagenized α-syn and α-syn + *atfs-1(lf) C. elegans* (Figure 2E).

### Retest: Neurodegeneration analysis of homozygous mutants

Following completion of the initial screen, 63 mutant lines were maintained that displayed the suppressor phenotype. Since our primary screen was performed with a 25X setting on a stereo fluorescent microscope, where we selected mutant lines that exhibited reduced numbers of DA neuron cell bodies compared to control *C. elegans,* our ability to accurately score DA neurodegeneration in detail remained incomplete. Therefore, all 63 candidate mutant lines were re-examined using compound fluorescent microscopy at the F6 generation (Figure 3A). Here, a more rigorous DA neurodegeneration analysis included a thorough examination of each of the six anterior DA neurons (4 CEPs and 2 ADEs) for intact cell bodies, as well as neuronal projections – where missing, degenerative, and broken neurites could be distinguished between unmutagenized α-syn and α-syn + *atfs-1(lf) C. elegans* (Figure 3C vs. 3D). From our analysis of all 63 mutant lines, we eliminated 41 candidates because they did not display significantly greater neurodegeneration compared to unmutagenized α-syn + *atfs-1(lf)* animals. The remaining 22 lines were significantly more neurodegenerative than unmutagenized α-syn + *atfs-1(lf)* animals and were retained for further analyses. Notably, of these 22 mutant lines, 9 exhibited significantly more neurodegeneration than did in *C. elegans* expressing α-syn (alone) (Figure 3B).

**Figure 3.**
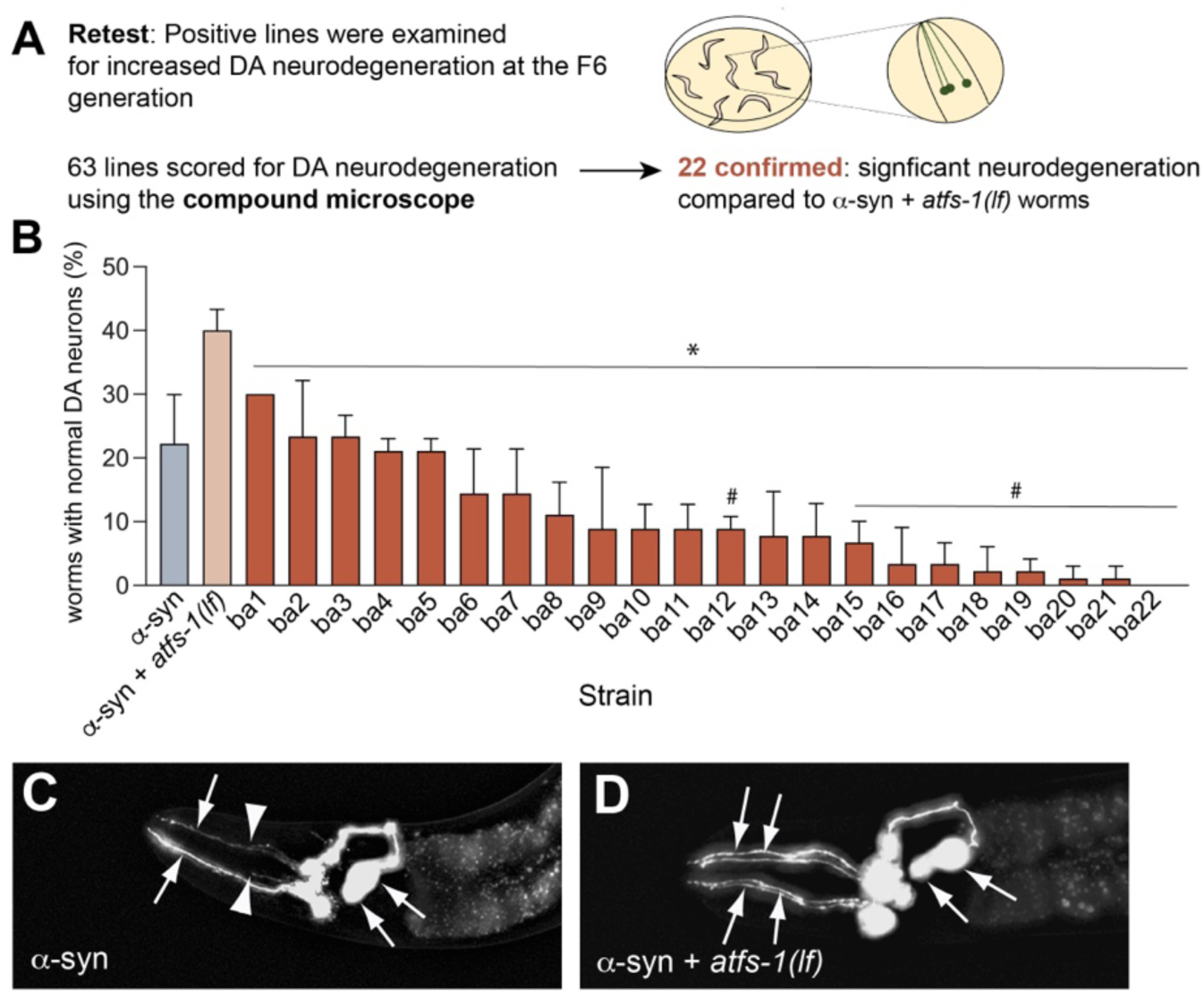
Retest for suppression of DA neuroprotection in α-syn + *atfs-1(lf)* worms using a compound fluorescent microscope. (A) Schematic of the screening protocol for retesting positive mutant lines. DA neurodegeneration (neurites and cell bodies) was assessed for each mutagenized line. (B) All 22 “*ba*” mutant lines, which have the α-syn + *atfs-1(lf)* genetic background, were identified from the screen as significantly suppressing DA neuroprotection compared to α-syn + *atfs-1(lf)* using Student’s *t*-tests p<0.05; N=3; n=30 (indicated with *). Additionally, 9 of the “*ba*” lines also significantly suppress DA neuroprotection compared to α-syn alone via Student’s *t*-tests (*ba12*, *ba15*-*ba22*); p<0.05; N=3; n=30 (indicated with #). Screening was performed on day 7 post-hatching. (C, D) These representative images portray the six anterior DA neurons in worms expressing GFP on day 7, post-hatching. Arrowheads indicate intact DA neurons and arrows indicate regions with degenerating and/or missing DA neurons. (C) This animal, which expresses GFP + α-syn in DA neurons, has four intact neurons (arrowheads) and two degenerating neurons (arrows). (D) This animal expresses α-syn + *atfs-1(lf)* + GFP and displays the full complement of 6 anterior DA neurons.

### Genomic sequencing and in silico comparative allelic analysis

All 22 mutant lines that consistently enhanced the α-syn-induced DA neurodegenerative phenotype and were continued for further analysis. To reduce the impact of nucleotide changes produced by EMS, we backcrossed the mutant lines to the original α-syn + *atfs-1(lf)* strain (UA460) three times. In all mutant lines, we confirmed significant neurodegeneration compared to the α-syn + *atfs-1(lf)* strain. All 22 mutant lines were then sequenced using the Illumina platform. The original *C. elegans* unmutagenized α-syn + *atfs-1(lf)* strain was sequenced using a combination of Illumina and Oxford Nanopore platforms. A reference genome was constructed from the α-syn + *atfs-1(lf)* strain for comparison to the mutants. The program GATK was used to identify SNPs within exons of the 22 mutant lines. In total, we performed ten rounds of EMS, thus it was possible that more than one mutant suppressor line could have originated from the same P0 animal. Thus, a script (GitHub) was created to compare the nonsense and missense SNPs within the mutant animals against each other. These analyses determined that the mutant lines harbored independent mutations. Genomic sequencing and analysis revealed that there were approximately three nonsense mutations and/or splice-site junction alterations per mutant line.

### Isolation of three null mutations that suppress α-syn + *atfs-1(lf)* neurodegeneration

To uncover the identity of the mutations, we adopted an RNAi knockdown approach of candidate genes that were mutated in our alleles. To facilitate this, we constructed a new *C. elegans* strain that is an RNAi-sensitive version of our screening background strain, UA460 [P*_dat-1_*::α-syn; P*_dat-1_*::GFP; *atfs-1*(*gk3094*)]. We crossed UA460 worms with a strain that enables DA neuron-specific RNAi knockdown. In these animals a mutation in the *sid-1* gene blocks systemic RNAi everywhere except the DA neurons, where it is rescued by virtue of *sid-1* expression driven from the P*_dat-1_* promoter. This cross created the final strain UA469 [*atfs-1*(*gk3094*); P*_dat-1_*::α-syn; P*_dat-1_*::GFP; *sid-1*(*pk3321*); P*_dat-1_*::*sid-1*; P*_myo-2_*::mCherry]. To connect alleles with potential genetic loci, we started by examining the nonsense mutations by RNAi knockdown of the corresponding genes. We determined significantly increased DA neurodegeneration was reproducibly observable in the UA469 background following RNAi for three genes. This method resulted in the assignment of these causal alleles: *ba5* to *jmjd-3.1*, *ba13* to *jmjd-1.2* (Figure 4 and Figure 5), and *ba20* to *twk-14* (Figure 4 and Figure 6). The newly described strain names are listed in Table 1.

**Figure 4.**
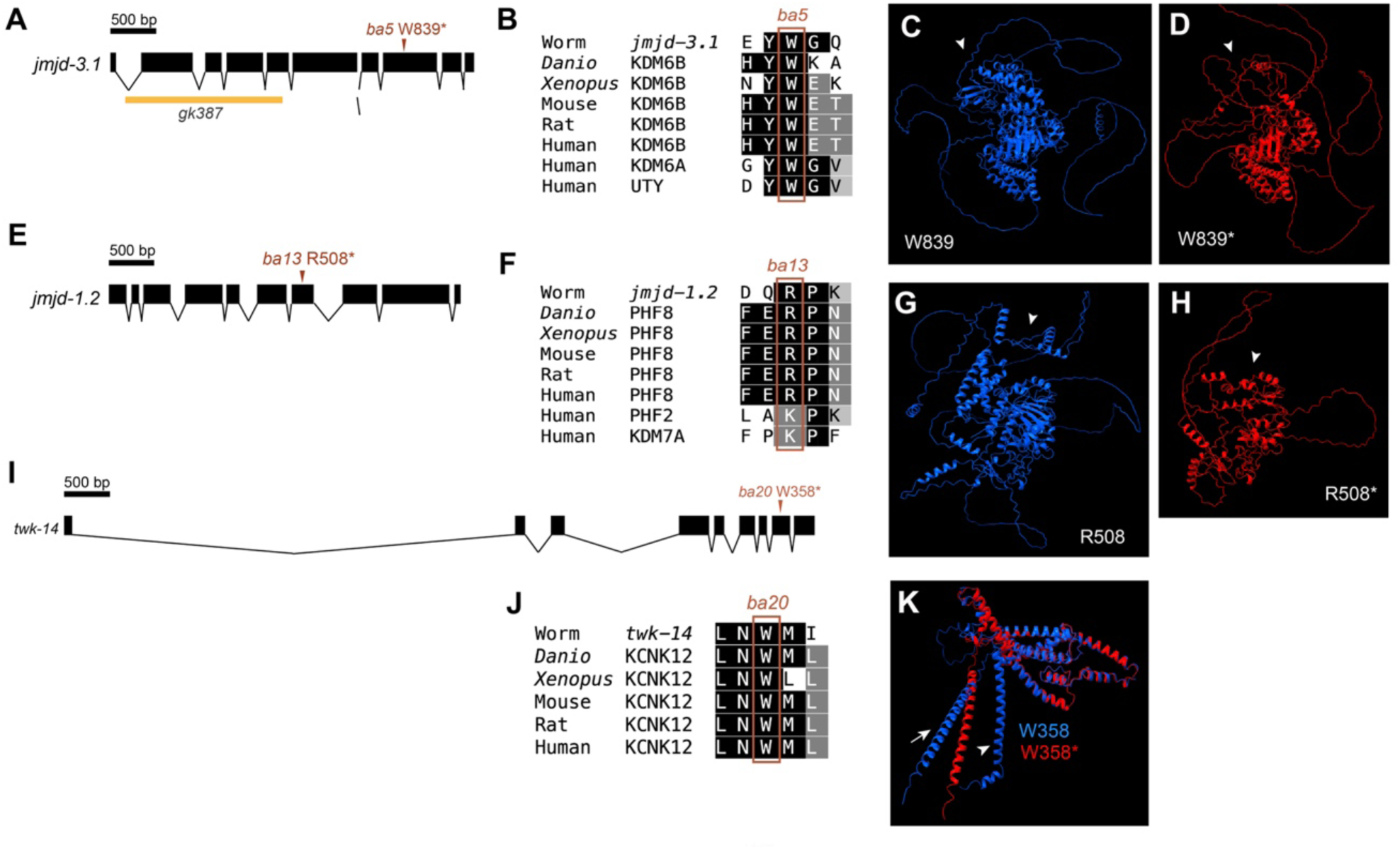
Three null mutations identified as modifiers of neurodegeneration in α-syn + *atfs-1(lf)* animals. (A) Gene structure of *jmjd-3.1* and position of mutant nucleotide in *ba5,* as verified by sequencing where a G → A mutation at nucleotide 2,517 from the ATG start that causes tryptophan to convert to a nonsense codon (W839*). (B) Alignment of amino acids adjacent to *jmjd-3.1(ba5)*, demonstrating conservation among species. (C, D) Three-dimensional structural models of the wildtype and mutant *jmd-3.1* were generated using the AlphaFold 3 server (Abramson et al., 2024). The resulting predicted structures were imported into ChimeraX for comparative analysis (Pettersen et al., 2021). The wildtype and nonsense mutant was superimposed using the Matchmaker algorithm within ChimeraX but are displayed as separate images for visual clarity. (E) Gene structure of *jmjd-1.2* and position of mutant nucleotide in *ba13*, as verified by sequencing where a C → T mutation at nucleotide 1,522 from the ATG start that causes conversion of an arginine to a nonsense codon (Arg508*). (F) Alignment of amino acids adjacent to *jmjd-1.2(ba13)*, demonstrating conservation among species. (G, H) Three-dimensional structural models of the wildtype and mutant *jmjd-1.2(ba13)* were generated as described in (C, D). (I) Gene structure of *twk-14* and position of mutant nucleotide in *ba20*, as verified by sequencing where a C → T mutation at nucleotide 1,074 from the ATG start that causes tryptophan to become a nonsense codon (Trp358*). (J) Alignment of amino acids adjacent to *twk-14(ba20)*, demonstrating conservation among species. (K) A three-dimensional structural model of both the wild-type and nonsense mutant proteins encoded by *twk-14* was generated as described in (C, D) except that the wild-type and nonsense structures of TWK-14 are presented as a single superimposed overlay to highlight their overlapping topologies.

**Figure 5.**
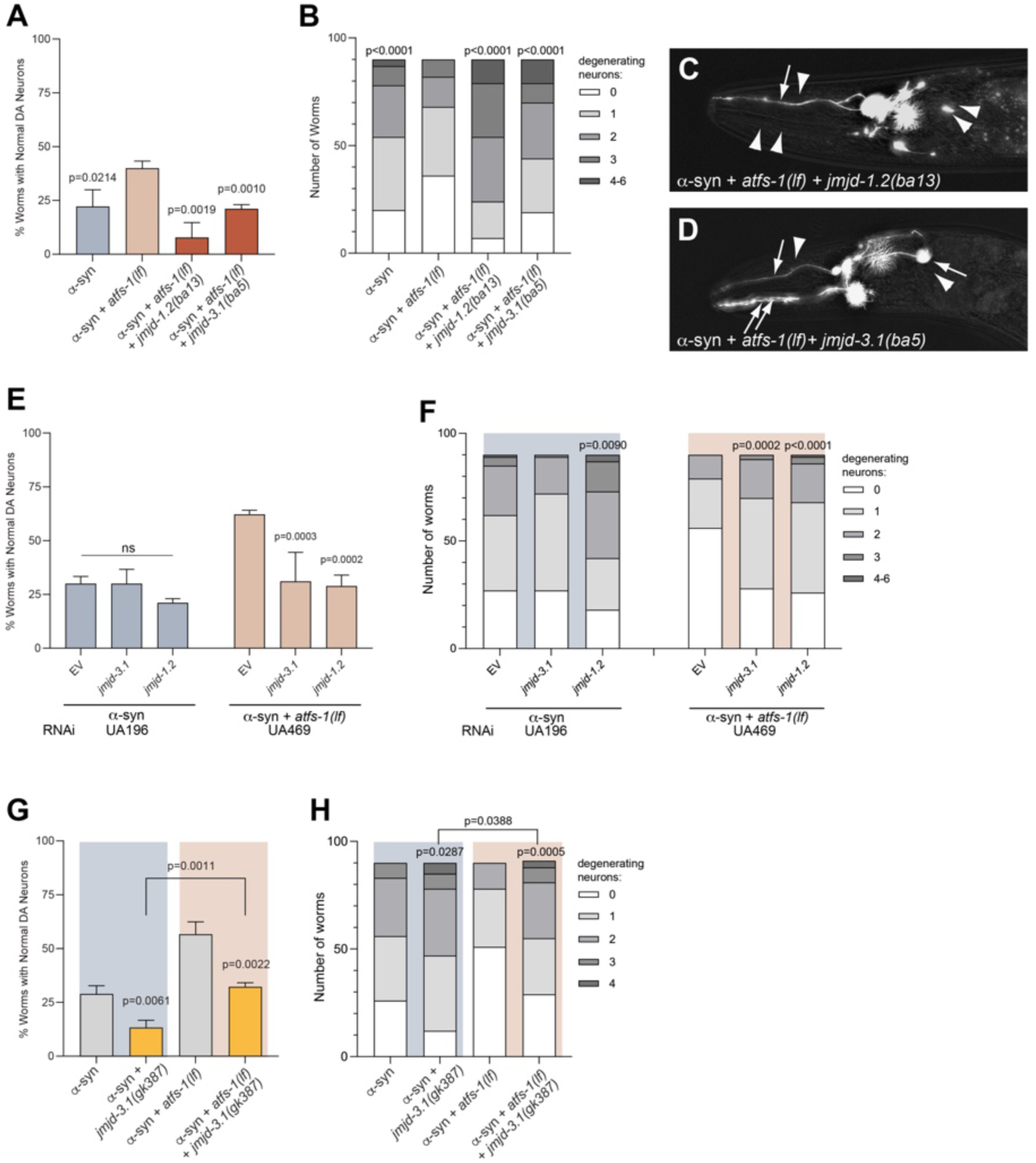
Characterization of *jmjd-1.2* and *jmjd-3.1* alleles in α-syn-induced neurodegeneration. (A) Mutant alleles identified from the screen (*ba13, ba5*) scored for DA neurodegeneration on day 7 post-hatching and compared to α-syn + *atfs-1(lf)* animals. Values represent mean + S.D. N=3; n=30; Student’s two-tailed type 2 *t*-test. (B) Data displayed in A were re-analyzed to examine the distribution of the entire population of neurons based on the indicated number of degenerating DA neurons (0 through 6) using a Chi square test in comparison to the α-syn + *atfs-1(lf)* control. N = 3; n = 30; *p* < 0.0001. (C, D) These images characterize the six anterior DA neurons in the mutant worms expressing GFP on day 7 post-hatching. Arrowheads indicate intact DA neurons and arrows indicate regions with degenerating and/or missing DA neurons. (C) An example *jmjd-1.2(ba13)* + α-syn + *atfs-1(lf)* animal with DA neurons illuminated by GFP. It has one intact (arrowhead) and five missing DA neurons (arrows). (D) This *jmjd-3.1(ba5)* + α-syn + *atfs-1(lf)* animal displays 4 intact anterior DA neurons illuminated with GFP (arrowheads). (E) RNAi knockdown of *jmjd-3.1* and *jmjd-1.2.* RNAi was performed in DA neuron RNAi-sensitive α-syn model strain (UA196) shown in blue and in DA neuron RNAi-sensitive α-syn + *atfs-1* model strain (UA469) shown in orange. RNAi-treated animals were scored for DA degeneration on day 7 post-hatching. EV=empty vector RNAi-treated control worms. Gene transcripts knocked down are specified on the x axis. Values represent mean ± S.D. (N=3; n=30; Student’s two-tailed type 2 *t*-test). (F) Data displayed in E were re-analyzed to examine the distribution of the entire population of neurons based on the indicated number of degenerating DA neurons (0 through 6) using a Chi square analysis test. N = 3; n = 30. (G) The *jmjd-3.1(gk387)* loss-of-function allele was crossed to both α-syn and α-syn + *atfs-1(lf)* animals and neurodegeneration scored on day 7 post-hatching. Values represent mean ± S.D. (N=3; n=30; Student’s two-tailed type 2 *t*-test). (H) Data displayed in G were re-analyzed to examine the distribution of the entire population of neurons based on the indicated number of degenerating DA neurons (0 through 6) using a Chi square analysis test. N = 3; n = 30.

**Figure 6.**
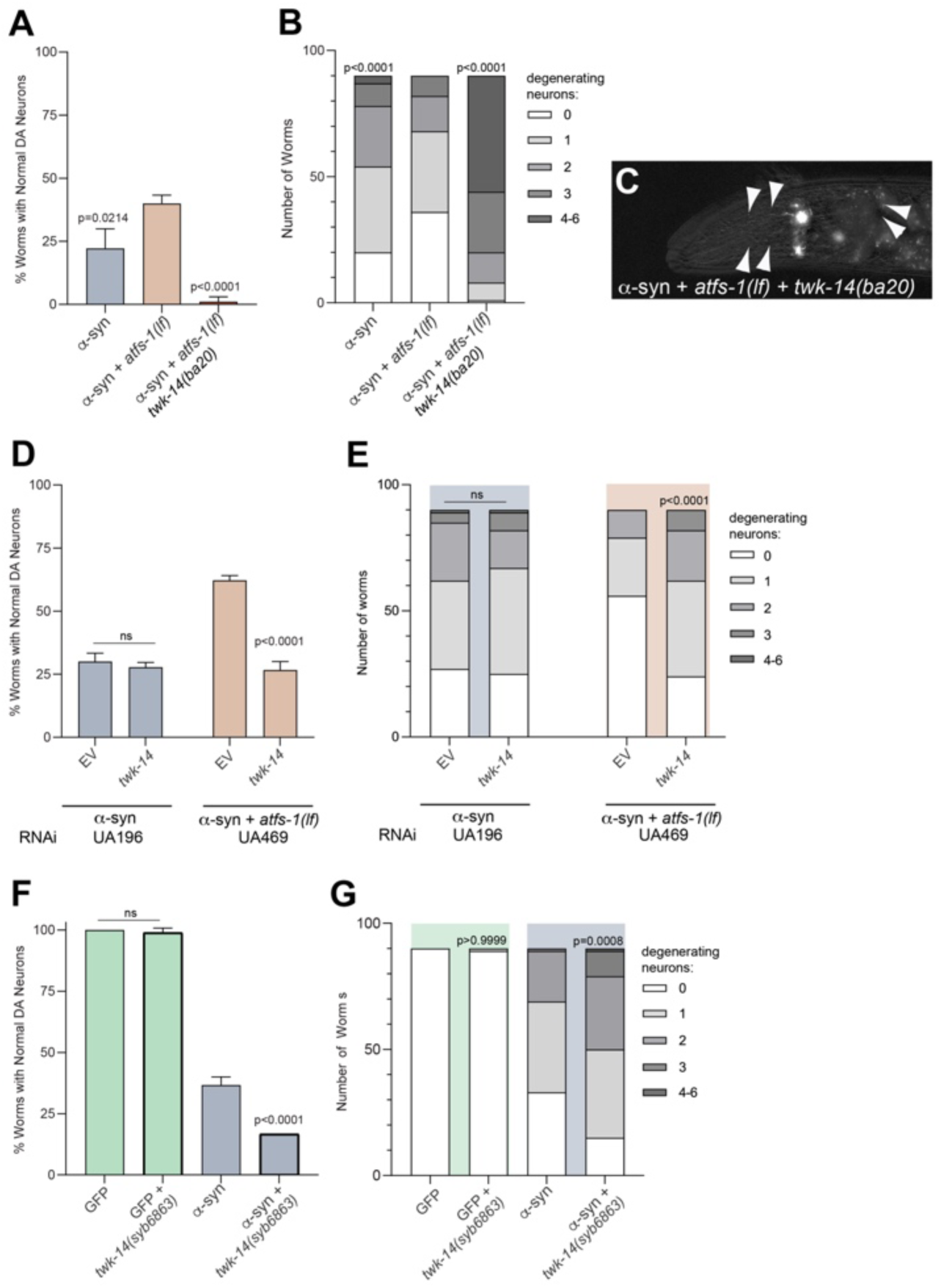
Characterization of a *twk-14* allele in α-syn-induced neurodegeneration. (A) The *twk-14* mutant strain (*ba20*) identified from the screen was scored for DA neurodegeneration on day 7 post-hatching and compared to α-syn + *atfs-1(lf)* animals. Values represent mean + S.D. N=3; n=30; Student’s two-tailed type 2 *t*-test. (B) Data displayed in A were re-analyzed to examine the distribution of the entire population of neurons based on the indicated number of degenerating DA neurons (0 through 6) using a Chi square test compared to the α-syn + *atfs-1(lf)* control. N = 3; n = 30; *p* < 0.0001. (C) This image displays the six anterior DA neurons in a *twk-14 (ba20)* mutant worm expressing GFP in the *afts-1(lf)* + α-syn background on day 7 post-hatching. Arrowheads indicate intact DA neurons and arrows indicate regions with degenerating and/or missing DA neurons. (D) RNAi was performed in DA neuron RNAi- sensitive α-syn model strain (UA196), shown in blue, and in DA neuron RNAi-sensitive α-syn + *atfs-1* model strain (UA469) in orange, both with DA neurons illuminated by GFP. Values represent mean ± S.D. (N=3; n=30; Student’s two-tailed type 2 *t*-test). (E) Data displayed in E were re-analyzed to examine the distribution of the entire neuron population based on the indicated number of degenerating DA neurons (0 through 6) using a Chi square test N = 3; n = 30. RNAi knockdown of *twk-14* in the UA196 strain did not lead to significant neurodegeneration. (F) Comparison of animals exhibiting DA neurodegeneration between worms expressing only GFP or GFP + CRISPR-edited *twk-14(syb6863)* shown in green, and these same animals crossed to worms with α-syn overexpressed in DA neurons, in blue. DA neurodegeneration assays performed on day 7 post-hatching. Values represent mean + S.D. N=3; n=30; Student’s two-tailed type 2 *t*-test. (G) Data displayed in F were re-analyzed to examine the distribution of the entire population of neurons based on the indicated number of degenerating DA neurons (0 through 6) using a Chi square test N = 3; n = 30.

### Loss of histone demethylases *jmjd-3.1* and *jmjd-1.2* cause significant neurodegeneration in α-syn *+ atfs-1(lf)* worms

Mutations in the Jumonji C domain lysine demethylase genes, *jmjd-3.1(ba5)* and *jmjd-1.2(ba13)*, were identified from our screen. The functional gene products encoded by both *jmjd-3.1* and *jmjd-1.2* have been reported to be required to establish or maintain an open chromatin state that allows ATFS-1 to bind and initiate transcription (Nargund *et al*. 2015; Merkwirth *et al*. 2016; Tian *et al*. 2016). The mutations in *jmjd-3.1(ba5)* and *jmjd-1.2(ba13)*, which were originally identified by Illumina sequencing, were confirmed using Sanger sequencing. The isolation of *jmjd-3.1(ba5)* and *jmjd-1.2(ba13)* represented strong internal validation of our screening strategy.

The *jmjd-3.1* gene contains ten exons and encodes a protein with 1,061 amino acids. The mutation in *ba5* generates a G → A transition mutation within exon 8 that results in a stop codon at amino acid 839, instead of a tryptophan (W839*) (Figure 4A). This residue is highly conserved across species (Figure 4B). We used AlphaFold3 to generate structural predictions of the W839* mutation and the wildtype variants. ChimeraX was then employed to reveal differences in hydrogen bonding predicted to alter protein folding between protein pairs resulting from the full-length wildtype protein (blue) and the nonsense mutant (red) (Figure 4C and 4D). Similarly, the *jmjd-1.2* gene contains ten exons and a protein consisting of 929 amino acids is encoded. The mutation in *ba13* represents a C → T transition within exon 7, whereby a stop codon results at amino acid 508, instead of an arginine (R508*) (Figure 4E). This basic residue is also highly conserved across species (Figure 4F). AlphaFold3-generated structural predictions, processed using ChimeraX to uncover protein folding differences between the full-length wildtype JMJD-1.2 protein (blue) and truncated R508* nonsense mutant variant (red), are depicted in Figure 4 (4G and 4H).

Quantitative analysis of DAergic neurodegeneration revealed that populations of both *jmjd-3.1(ba5)* and *jmjd-1.2(ba13)* displayed significantly more neurodegeneration than did unmutagenized α-syn + *atfs-1(lf)* worms (Figure 5A). Similarly, when we examined the severity of all DA neurodegenerative changes within a population, depicted as neuron counts representing the distribution of degenerating DA neurons, both *jmjd-3.1(ba5)* and *jmjd-1.2(ba13)* displayed significantly more DA neurodegeneration than unmutagenized α-syn + *atfs-1(lf)* worms (Figure 5B). *jmjd-1.2(ba13)* also displayed visibly more severe neuronal deficits compared to *jmjd-3.1(ba5)* (Figure 5C and 5D).

DA-sensitive RNAi knockdown within worms expressing α-syn + *atfs-1(lf)* (strain UA469) was used in our assignment of causal alleles *jmjd-3.1(ba5)* and *jmjd-1.2(ba13)*. In this background, knockdown of either gene compared to empty vector (EV) RNAi-treated controls resulted in significantly more worm populations with DA neurodegeneration (Figure 5E, orange bars). Likewise, the distribution of degenerating DA neurons displayed significantly more severe neurodegenerative changes within the population of neurons from both *jmjd-3.1* and *jmjd-1.2,* compared to EV control in the α-syn + *atfs-1(lf)* RNAi worms (Figure 5F, orange); this suggested that the enhanced α-syn neurodegeneration observed might require the *atfs-1(lf)* background. To test whether the neurodegenerative phenotype was independent of *atfs-1(lf),* we employed a DA neuron-sensitive RNAi strain of worms expressing of α-syn (strain UA196), this time in an *atfs-1* wildtype background; no significant difference in neurodegeneration between the EV control, *jmjd-3.1,* or *jmjd-1.2* RNAi was observed in worm populations (Figure 5E, blue bars). However, a re-analysis of these data based on neuron counts revealed a subtle distinction in that *jmjd-1.2* RNAi, but not *jmjd-3.1,* exhibited a significant enhancement in the distribution of DA neurons with more degeneration (Figure 5F). We attribute this distinction to the expression pattern of the gene transcripts, where *jmjd-1.2* is expressed in all DA neurons, whereas *jmjd-3.1* expression is more limited (Taylor et al., 2021).

To further characterize *jmjd-3.1*, we obtained an existing allele [*jmjd-3.1(gk387)*] with a deletion that removes 1,600 bp corresponding to exons 2-4 (Figure 4A); it results in a functional null (Agger *et al*., 2007). We crossed *jmjd-3.1(gk387)* to α-syn + *atfs-1(lf)* and to α-syn alone to discern whether the neurodegeneration observed is dependent on an absence of *atfs-1* activity. We found significantly more neurodegeneration when the *jmjd-3.1(gk387)* null allele was crossed to both α-syn + *atfs-1(lf)* and to α-syn alone (Figure 5G and Figure H). Taken together, we conclude that reduction of *jmjd-3.1* function influences α-syn-induced neurodegeneration in a manner that is independent of *afts-1*. Moreover, in consideration of the *jmjd-3.1* RNAi data (Figure 5E), this effect appears to occur cell non-autonomously.

While we did not examine a *jmjd-1.2* mutant within this same context, the presence of both JMJD proteins is known to be required for an open chromatin state, and for the ATFS-1-dependent expression of UPR^mt^ genes (Nargund *et al*. 2015; Merkwirth *et al*. 2016; Tian *et al*. 2016; Muñoz-Carvajal and Sanhueza, 2020). However, chromatin remodeling can also occur independently of ATFS-1. Activation of the UPR^mt^ requires the di-methylation of histone H3K9 by histone methyltransferase MET-2 and LIN-65, a nuclear co-factor. For example, in the absence of ATFS-1 significantly more LIN-65 accumulates in the nucleus; perhaps emphasizing that increased mitochondrial stress stimulates LIN-65 trafficking (Tian *et al*. 2016). While mechanisms for how the Jumonji methyltransferases are stimulated by mitochondrial stress remain unresolved, both JMJD-1.2 and JMJD-3.1 require alpha-ketoglutarate and iron, which are affected by mitochondrial dysfunction (Merkwirth *et al*., 2016). Notably, it was recently reported that *C. elegans* fed a diet depleted of iron displayed an enhanced UPR^mt^ response (Das et al., 2025). We also previously reported that age-dependent α-syn-induced degeneration of DA neurons was partially rescued by the iron chelator desferoxamine (Patel et al., 2018). These results collectively indicate that epigenetic regulation of the UPR^mt^ involves coordination of both endogenous and exogenous signals to maintain proteostasis and neuronal survival.

### Loss of *twk-14* causes significantly enhanced neurodegeneration in α-syn *+ atfs-1(lf)* worms

Our screen also identified an allele of *twk-14(ba20).* TWK-14 is a two-pore domain potassium leak channel (K2P) that is homologous to human KCNK12. The mutation isolated in *twk-14(ba20)* was initially identified by Illumina sequencing and the genetic lesion was confirmed by Sanger sequencing. The *twk-14* gene contains nine exons and encodes a protein with 463 amino acids. The mutation in *ba20* represents a C → T transition mutation within exon 8 that results in a stop codon at amino acid position 358 instead of a tryptophan (W358*) (Figure 4I). This residue is highly conserved across species (Figure 4J). To visualize the potential impact of this change, structural predictions of W358* and wildtype TWK-14 were generated with AlphaFold3. Protein folding differences were revealed by superimposing the nonsense mutant (red) and the full-length wildtype protein (blue) using ChimeraX. The resulting image identified a missing C-terminal alpha-helical region (arrowhead – on WT) and a shifted alpha-helix (arrow – on WT) (Figure 4K).

We performed an in-depth analysis of DA neurodegeneration with the *twk-14(ba20)* allele within the α-syn + *atfs-1(lf)* background (Figure 6A-6C). Populations of *twk-14(ba20) C. elegans* displayed significantly more DA neurodegeneration than unmutagenized α-syn + *atfs-1(lf)* animals (Figure 6A). Similarly, the severity of degeneration assessed by the distribution of DA neurons counted in *twk-14(ba20)* animals was greater in comparison to unmutagenized α-syn + *atfs-1(lf)* worms (Figure 6B).

To initially assign the causal allele, *twk-14*(*ba20*), a DA neuron-sensitive RNAi strain expressing α-syn + GFP + *atfs-1(lf)* (strain UA469) was used for *twk-14* knockdown. In the absence of *atfs-1*, RNAi knockdown of *twk-14* resulted in significantly more animals with DA neurodegeneration when compared to EV RNAi-treated controls (Figure 5D, orange bars). We also determined that significantly more severe degeneration occurred within populations of DA neurons counted following *twk-14* RNAi compared to EV control in α-syn + *atfs-1(lf)* RNAi worms (Figure 5E, orange). To examine whether this enhanced neurodegenerative phenotype was dependent on *atfs-1(lf)*, we knocked down *twk-14* in a DA-sensitive RNAi strain of worms overexpressing α-syn (and GFP), but with wildtype *atfs-1* (strain UA196). No significant difference in neurodegeneration between the EV control and *twk-14* RNAi was observed (Figure 5D, blue bars). The distribution of individual degenerated neurons between EV and *twk-14* RNAi-treated animals were also not different in this background (Figure 5E, blue). Taken together, these data suggest that *atfs-1* loss-of-function might be required to observe enhanced α-syn neurodegeneration.

Since the identification of *twk-14* as a modifier of neurodegeneration is a novel finding, we wanted to independently confirm this genetic lesion by additional means. We used CRISPR technology to edit the chromosomal *C. elegans twk-14* gene and recapitulate the isolated *ba20* allele through genome engineering. A strain was created where a SNP was introduced into the endogenous *twk-14* locus that encoded a protein variant reflective of the W358* amino acid nonsense mutation in *twk-14(ba20)* mutant animals (Figure 4I). We crossed these edited animals to worms expressing GFP alone (BY250) in the DA neurons to obtain strain UA477 {*twk-14(syb6863)*; [*vtIs7*] P*_dat-1_*::GFP}. We learned that *twk-14(syb6863)* animals do not independently display DA neurodegeneration at either the worm or neuron population levels (Figure 6F, green; Figure 6G, green). We next crossed the *twk-14(syb6863)* CRISPR mutant strain with worms expressing DAergic α-syn + GFP and obtained strain UA478 {*twk-14(syb6863)*; [*baIs11*] P*_dat-1_*::α-syn; P*_dat-1_*::GFP}. These animals exhibited significantly increased neurodegeneration compared to α-syn worms, both when scored at the level of whole animal populations (Figure 6F, blue) and when the distribution of neurons within populations was quantified (Figure 6G, blue). Unfortunately, repeated attempts to cross *twk-14(syb6863)* with α-syn + *atfs-1(lf)* worms were unsuccessful, likely because *twk-14* and *atfs-1* are genetically linked and only <2.1 cM apart, on chromosome V.

The outcomes of this analysis indicated that although the CRISPR-generated *twk-14(syb6863)* allele successfully phenocopied the neurodegeneration of the isolated *ba20* allele, there remained a distinction between these genetic variants and the results of the *twk-14* RNAi knockdown (Figure 5G, 5H vs. 5I, 5J). Importantly, based on the complete neuronal connectivity maps (wormatlas.org) and the comprehensive gene expression profile of all individual neurons (Taylor et al., 2021), *twk-14* is not expressed in DA neurons. Rather, it is expressed within select sensory neurons in the head, where its limited localization includes the neuropeptide-secreting FLP neurons, postsynaptic to the DA neurons in a defined neuroregulatory circuit. Therefore, since the RNAi experiments are specific to DA neuron knockdown, we conclude that the effect observed in *twk-14* genetic mutants likely occurs cell non-autonomously.

The human genome encodes fifteen K2P proteins, subclassified into six subfamilies based on pharmacological and biophysical properties (Schreiber and Seebohm, 2026). Well-studied K2P subfamily members have measurable current at the plasma membrane and are associated with human pathologies, including migraine with aura (Lafrenière *et al*., 2010), cardiac arrhythmias (Chen *et al*., 2014), and cancers (Williams *et al*., 2013). The K2P channel family that KCNK12 is part of is the THIK (tandem pore domain <u>h</u>alothane inhibited <u>K</u>^+^ channel) subfamily. Studies on THIK family member activity were performed on the rat homologs, referred to as THIK1 (KCNK13) and THIK2 (KCNK12). THIK1/KCNK13 is expressed ubiquitously and produces background K+ currents, whereas THIK2/KCNK12 is silent and limited in expression to the brain and kidney (Rajan *et al*., 2001). It is intriguing that KCNK12 is inactive, considering it is 62% identical to KCNK13. However, the N-terminal domain of THIK2/KCNK12 has an arginine-rich motif (RRSRRR) that acts as an ER retention/retrieval signal motif, lowering localization of this channel at the plasma membrane (Chatelain *et al*., 2013; Blin *et al*., 2014; Renigunta *et al*. 2006).

Structure-function analyses have determined THIK1 and THIK2 form heterodimers that distribute to the plasma membrane, with measurable current (Chatelain *et al*., 2013, Blin et al., 2014). Natively, however, THIK2/KCNK12 is a homodimer at the ER, with low intrinsic activity (Theilig et al., 2008; Bichet et al., 2015). Most research on KCNK12 has focused on its cellular localization and/or heterologous expression studies on ion channel activity (Aggarwal *et al*., 2021). It is possible that the *C. elegans* KCNK12 homolog, TWK-14, localizes to the ER as a homodimer, and/or has an unknown heterodimeric partner. While the human genome encodes 15 K2Ps, there are 47 K2Ps predicted with in the *C. elegans* genome that are largely uncharacterized (Hobert, 2013; Zhou *et al*., 2022).

ATFS-1 regulates the UPR^mt^ in *C. elegans*, where it normally acts as a safeguard against transient stressors affecting mitochondrial function (Durieux *et al*., 2011; Rauthan *et al*., 2013; Pellegrino *et al*., 2014). However, this protective cellular program turns toxic when sustained stress, such as that imposed by mitochondrial DNA damage or other genetic perturbations, results in constitutive UPR^mt^ activation (Melber and Haynes, 2018). We previously reported that co-expression of α-syn and ATFS-1 synergistically potentiated DA neurotoxicity in *C. elegans* (Martinez *et al*., 2017). Importantly, this neurotoxicity was attenuated in an *atfs-1(lf)* background where UPR^mt^ activation was blocked (Figures 1A, 3C, 3D). Conversely, α-syn-induced DA neurodegeneration is increased in *atfs-1* gain-of-function *(gf)* animals (Martinez *et al*., 2017). These effects on DA neuron survival do not occur without α-syn in the *atfs-1*(*lf*) or *(gf)* backgrounds, suggesting that neuroprotection from α-syn-induced neurodegeneration in *atfs-1(lf)* occurs through a UPR^mt^ compensatory mechanism (Martinez *et al*., 2017). However, it is unclear what molecular components are associated with *atfs-1(lf)* in α-syn-expressing DA neurons.

These data provided a background through which a classical forward genetic screen was performed to identify molecular components linked to either the UPR^mt^ or α-syn neuroprotection (or both) in DA neurons. However, it is unclear what molecular components are associated with *atfs-1(lf)* in α-syn-expressing DA neurons. We were interested in identifying molecular components that, when mutant, lead to increased DA neuronal degeneration from UPR^mt^ overactivation, with the idea that these represented inherently neuroprotective factors. Using *C. elegans* expressing α-syn specifically in the DA neurons that display significant neuroprotection when crossed with a deletion mutation in *atfs-1* (Figure 1B), we identified a suite of genetic modifiers. Among these, we isolated mutations in two paralogous genes, *jmjd-3.1(ba5)* and *jmjd-1.2(ba13)*. Previous studies had established that the products of these Jumonji C domain lysine demethylase genes were important for transcriptional regulation of ATFS-1.

Strikingly, a recent report wherein the neurotoxins MPTP and 6-OHDA were used to induce DA neurodegeneration in cell cultures from mice demonstrated that JMJD3 demethylase activity works in a protective capacity to decelerate PD-like progression; clearance of the epigenetic mark, H3K27me3, by JMJD3 on the promoter of the SNAI2 transcription factor stimulates upregulation the potent neuroprotective enzyme, HIF1α (Dong and Guo, 2023). It is yet to be discerned if this response to acute neurotoxins is recapitulated when neurons are under chronic UPR^mt^ activation in mammalian α-syn models of PD. Nevertheless, our data on *jmjd-3.1* and *jmjd-1.2* with the transgenic *C. elegans* α-syn model support a conserved neuroprotective function for Jumonji demethylases in PD.

Additionally, we identified a mutant allele *(ba20)* of the *twk-14* gene that encodes a conserved neuromodulator of the K2P channel family. KCNK12, the human ortholog of TWK-14, is classified as understudied and is a member of a “druggable” family by multiple functional genomics consortia (Koscielny et al., 2017; Oprea et al., 2018; Kustatscher et al., 2022). Currently, we are unaware of any reports describing a role for any K2P channel in modulating the UPR^mt^ and/or neurodegeneration. In this regard, our screen suggests a new role for K2P channels in general, and TWK-14, specifically.

In summary, experimental strategies to target the pivotal balance in proteostasis that must be maintained to avoid both intrinsic genetic, as well as epigenetic and environmental challenges to DA neuron survival, represent a substantive opportunity for discovery. The outcomes of this screen, and continued investigation of mechanistic insights unearthed by these worms, exemplify the utility of *C. elegans* as a preclinical model with potential to reveal previously unforeseen therapeutic paths.

## Acknowledgements

The authors would like to thank Dr. Cole Haynes (University of Massachusetts Medical School) for insightful discussions. We are grateful to Dr. Janna Fierst (Florida International University) for generously providing access to the ONT GridION, and Joshua Millwood for his expertise and assistance extracting high molecular weight DNA and library preparation for ONT sequencing. We also thank Dr. Randy Blakely (Florida Atlantic University) for sharing strain BY250. Some strains were provided by the CGC, which is funded by NIH Office of Research Infrastructure Programs (P40 OD010440). This research was supported by grants from the National Institutes of Health (R15NS106460 and R03TR004460) to KAC.

